# Obesity-Associated Alterations in Lung Function in Mice Measured with Head-Out Plethysmography

**DOI:** 10.1101/2022.05.25.493464

**Authors:** Stephanie M Bruggink, William P Pederson, Kyle P Kentch, Jason M Kronenfeld, Julie G Ledford, Benjamin J Renquist

**Affiliations:** Physiological Sciences GIDP, University of Arizona, Tucson, AZ 85724; Animal and Comparative Biomedical Sciences, University of Arizona, Tucson, AZ 85721; Cellular and Molecular Medicine, University of Arizona, Tucson, 85724

**Keywords:** Obesity, Head-Out Plethysmography, Forced Oscillation Technique, Hepatic Steatosis, Mice

## Abstract

Declines in lung function worsen quality of life and increase the risk of mortality. Obesity and non-alcoholic fatty liver disease are associated with worsened lung function. To investigate this association, we assessed lung function in lean and diet-induced obese conscious mice using our newly developed leak-free head-out plethysmography system. Obesity was associated with increased volume (P<0.0001), minute ventilation (volume per minute; P<0.0001), mid-expiratory flow (flow rate at 50% expiratory volume; P<0.0001), end-inspiratory pause (pause at end of inspiration; P<0.0001) and decreased expiratory time (P<0.0001). We next compared the response to methacholine (0, 25, 50, 100 mg/ml in PBS flow 0.2ml/30sec) measured using our head-out plethysmography system with forced oscillation technique (using the standard flexiVent system) measures taken in the same mice. Many of the measures gathered using head-out plethysmography were associated with measures collected using the forced oscillation technique. Minute ventilation was most significantly associated with maximal airway resistance, maximal airway elastance, tissue damping, and tissue elastance (r=-0.59 P<0.0001; r=-0.54 P<0.005; r=-0.48 P<0.005; r=-0.40 P<0.005 respectively). Volume, corrected for energy expenditure, was most significantly associated with maximal resistance of the conducting airways (r=-0.57 P<0.0001). Although fatty liver is associated with changes in lung function, we found neither hepatic vagotomy nor knocking down obesity-induced hepatic GABA production improved lung function in obese mice. Still, our head-out plethysmography system is ideal for assessing the response to interventions aimed at improving obesity-associated declines in lung function.

## Introduction

Decreases in lung function are associated with declines in both physical health and quality of life(1, 2). Obesity is associated with impairment in lung function, measured by decreases in both volume and flow(3–6). Obesity-associated impairments in lung function may increase the risk for development of lung disorders or exacerbate symptoms in diseases such as asthma(7, 8). A mechanistic understanding of obesity-associated decreases in lung function is key to developing effective treatments and preventative measures(9–11).

The current “gold-standard” method to assess lung function in mice is forced oscillation technique, which assesses lung mechanics by measuring impedance of airflow into the lung of an anesthetized, paralyzed, tracheostomized, and ventilated mouse(12). This technique is precise, assessing resistance and compliance of the lung, but, is terminal (unless mouse is orotracheally intubated) and cannot be applied to assess the voluntary control of breathing(13).

We have previously described the validation of a leak-free, head-out plethysmography system for use in mice(14). Head-out plethysmography assesses lung function by placing mice in an airtight chamber with their head protruding. Using a pressure transducer chamber pressure changes are measured as the ribcage expands and contracts with each breath. We improved the previously described systems to eliminate air leak around the neck of the mouse. We created an inflatable balloon cuff to gently inflate around the neck of the mouse without impacting breathing, allowing for accurate measures of volume and flow.

Mouse models of obesity recapitulate many of the features of metabolic syndrome including ectopic lipid accumulation. Lipid accumulation in the liver, hepatic steatosis, is associated with declines in lung function, including reduced forced expiratory volume over 1 second and forced vital capacity(15). Non-alcoholic fatty liver disease is also associated with lung disorders such as chronic obstructive pulmonary disorder, asthma, obstructive sleep apnea, and pulmonary cancer(16–18). We set out to investigate the association between hepatic steatosis and obesity-associated declines in lung function in mice.

We recently established that hepatic gamma aminobutyric acid (GABA) inhibits the hepatic vagal afferent nerve (HVAN) resulting in insulin resistance and hyperinsulinemia(19, 20). Knocking down GABA-transaminase (GABA-T) mRNA expression or pharmacologically inhibiting GABA-T decreases hepatic GABA production and limits obesity induced hyperinsulinemia and insulin resistance(20). We designed a study to assess the role of the hepatic vagal nerve and liver GABA production in obesity-induced changes in lung function. A subset of these results have previously been reported as an abstract(21).

## Materials & Methods

### Mice

Studies were approved by the Animal Use and Care Committee at the University of Arizona. Studies used male C57BL/J6 mice (Control diet: Tekland WI 7013 NIH-31). High fat diet (HFD: Tekland WI TD 06414) was provided at 12 weeks of age for 9 weeks. At 21 weeks of age, we assessed lung function in chow fed lean (24.4 + 0.7 grams) and diet-induced obese mice (40.8 + 1.2 grams) with head-out plethysmography and subsequently measured lung mechanics using forced oscillation technique in a subset of mice(14).

### Antisense Oligonucleotide & Hepatic Vagotomy Surgery

We treated diet-induced obese mice (6-8 month age) with a control antisense oligonucleotide (ASO; 12.5 mg/kg bi-weekly for 4 weeks) or an ASO targeted against GABA-Transaminase (ABAT ASO) to limit hepatic GABA production(19, 22). To assess the role of the hepatic vagal nerve, we performed sham or hepatic vagotomy surgeries in 12-week old male mice and assessed lung function at 0, 3, 6, and 9 weeks on HFD(22).

### Breathing Measures

For all studies, a 5-minute baseline measurement was collected, then mice were exposed to nebulized (Kent Scientific, Torrington, CT; Aeroneb lab nebulizer small unit #AG-AL1100 & Aeroneb Lab Control Module #AG-AL7000) methacholine (MP 220 Biomedicals, LLC; total of 0, 5, 10, and 20 mg aerosolized in PBS) onto the mouse’s head. Mice fully recovered in their home cage for at least 1 h between doses. Five-minute baseline measurements were averaged to calculate tidal breathing. Time course data is presented every 30 seconds as 1-minute rolling averages. When applicable, data is normalized to the average of the 5-minute baseline measurement.

In a subset of lean and obese mice, 24-hours after collecting breathing measurements using head-out plethysmography we performed forced oscillation technique. Mice were anesthetized (Urethane Sigma #U2500-500G; 1 g/kg) and paralyzed (Pancuronium bromide Sigma #P1918; 0.8 mg/kg at 10 μl per gram of body weight). The trachea were cannulated with a 19-guage cannula and connected to a computer-controlled piston-ventilator (flexiVent, SCIREQ Inc., Montreal, Qc, Canada) for mechanical ventilation. We used a script to determine the bronchoconstrictive response to methacholine (MP Biomedicals #190231; total 0.21, 1.75, 3.5, and 7 mg nebulized in PBS) with a deep inflation preceding exposure to methacholine (Kent Scientific, Torrington, CT; Aeroneb Lab Nebulizer Small Unit #AG-AL1100 & Aeroneb Lab Control Module #AG-AL7000).

### Statistics

Data analyses were performed in SAS (SAS Inst., Cary, NC). To determine association between forced oscillation technique and head-out plethysmography measures, we performed Pearson’s correlation analyses. To identify the maximal ability of the head-out plethysmography variables to predict measures from forced oscillation technique, we performed a linear regression allowing for the stepwise removal of variables with P > 0.1. Both regression and correlation analyses were performed with all mice and again with mice segregated by diet. A repeated measures mixed model ANOVA was used to assess effect of vagotomy and Student’s T-test to assess the effect of ABAT ASO. As applicable, Tukey’s adjustment was used to correct for multiple comparisons.

## Results

### Obesity-Associated Alterations in Breathing

Obesity was associated with significant alterations in tidal breathing; increases in volume (volume assessed from pressure change as the ribcage expands; 0.06 + 0.01 ml; P<0.0001), minute ventilation (volume in millimeters per minute; 18.5 + 2.6 ml/min; P<0.0001), expiratory time (−11.7 + 2.6 msec; P<0.0001), mid-expiratory flow (flow rate at 50% of expiratory volume; 0.8 + 0.2 ml/sec; P<0.0001), and end-inspiratory pause (pause occurring at the end of inspiration; 6.5 + 2.4 msec; P=0.009) (Figure 1). Weights are reported in online data Supplement Table E1.

**Figure 1.**
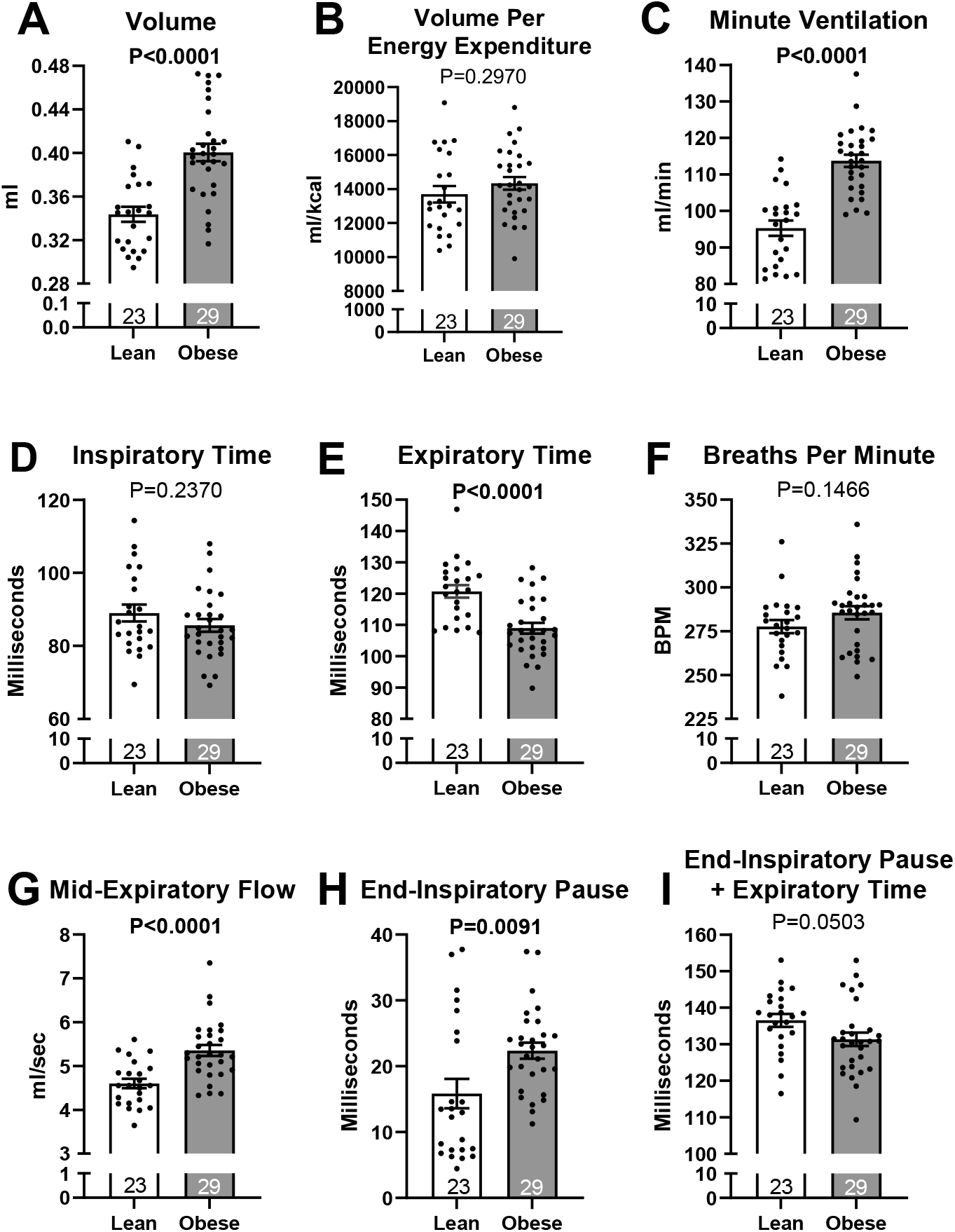
Tidal breathing comparison between lean and diet-induced obese male mice for variables collected using head-out plethysmography. The authors have reported a subset of group (n=9 lean mice tidal breathing & response to methacholine) previously(14). For all graphs lean n=23 obese n=29; results presented as mean + SEM. Analyzed with Student’s T-Test.

Lean and obese mice responded similarly to methacholine as measured using head-out plethysmography (Figure 2) and the forced oscillation technique (Figure 3). We have reported the data for a subset of the lean mice (9 lean mice tidal breathing and response to methacholine) previously(14).

**Figure 2.**
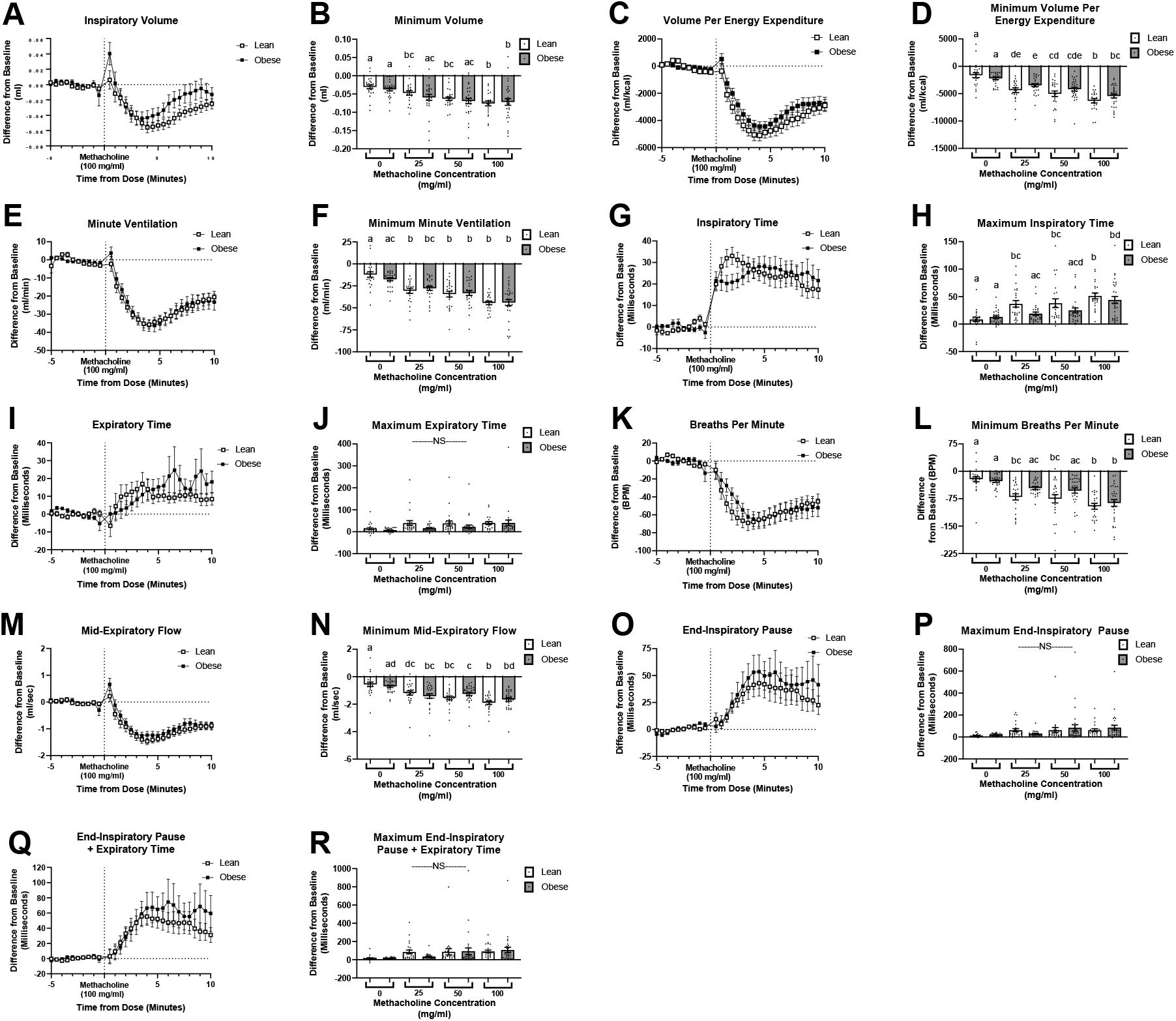
Response to methacholine in lean and diet-induced obese male mice. A, C, E, G, I, K, M, O, Q) Time course response to aerosolized 100 mg/ml of methacholine for 30 seconds at flow rate of 0.2 ml/30sec given at time 0. Data expressed as a difference from average 5-minute baseline measurement. B, D, F, H, J, L, N, P, R) Maximal change for each variable after exposure to different concentrations of methacholine (exposure 30 seconds flow rate of 0.2ml/30 sec). Different letters signify statistically significant difference between bars. The authors have reported a subset of group (n=9 lean mice tidal breathing and response to methacholine) previously(14). For all graphs lean n=23 obese n=29; results presented as mean + SEM. Analyzed with mixed model ANOVA.

**Figure 3.**
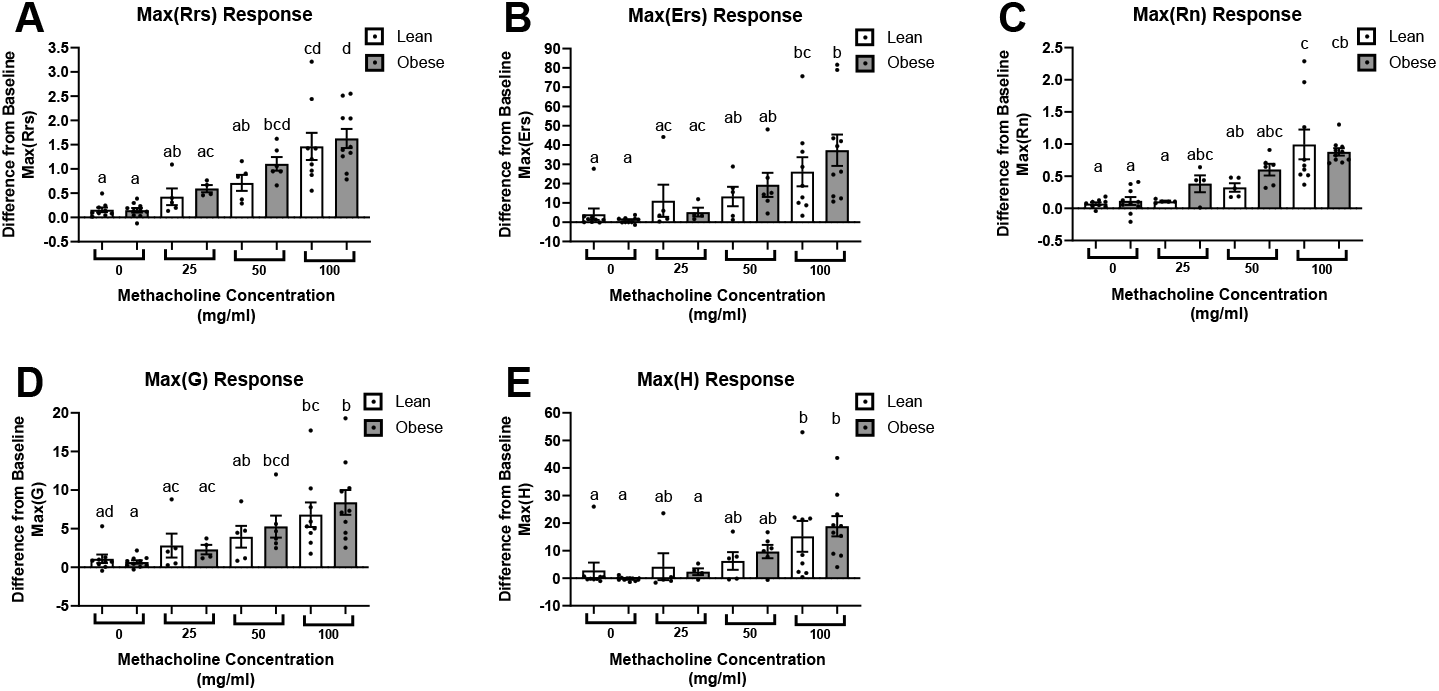
Response to methacholine in lean and diet-induced obese male mice measured using forced oscillation technique with the FlexiVent system. A) Maximal airway resistance, Max(Rrs), B) Maximal airway elastance, Max(Ers), C) Maximal resistance of conducting airways, Max(Rn), D) Maximal tissue damping, Max(G), and E) Maximal tissue elastance, Max(H). Results presented as maximal difference from baseline measurement (3 mg/ml methacholine flow 0.07ml/10sec). Mice were exposed to dose of methacholine for 10 seconds. Different letters correspond with statistically significant difference between bars. Lean n=9 obese n=10 results presented as mean + SEM. Analyzed with mixed model ANOVA.

### Head-out Plethysmography Measures Associated with Forced Oscillation Technique Measures

To compare the two systems, we calculated the maximal change from baseline for each variable at each dose of methacholine with the two systems. The response at 0, 25, 50, and 100 mg/mL flow 0.2ml/30sec measured using head-out plethysmography was associated with the equivalent dose of 3, 25, 50, and 100 mg/ml flow 0.07 ml/10sec collected using forced oscillation technique.

Variables collected with the two systems were correlated (Table 1-All; Figure 4). Minute ventilation 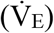 was the variable most significantly associated with maximal airway resistance (Max(Rrs) level of constriction of the lungs), maximal airway elastance (Max(Ers) elastic stiffness of respiratory system), maximal tissue damping (Max(G) resistance of peripheral airways and tissues – reflecting energy dissipation in alveoli), and maximal tissue elastance (Max(H) reflecting energy conservation in alveoli) (r=-0.59 P < 0.0001, r=-0.54 P=0.0004, r=-0.48 P=0.0001, and r=-0.40 P=0.0018 respectively). Volume accounting for energy expenditure was most significantly associated with maximal resistance of the conducting airways (Max(Rn); r=-0.57 P<0.0001). Regressions using head-out plethysmography variables explained up to 53% of the variation in forced oscillation technique variables (Table 2-All).

**Table 1.** Statistically Significant Regressions Comparing Maximal Response to Methacholine Measured via FlexiVent with Response to Methacholine Measured Via Head-Out Plethysmography in Order of Most Significant Association.

| Max (Rrs) |  |  | Max (Ers) |  |  | Max (Rn) |  |  | Max (G) |  |  | Max (H) |  |
| --- | --- | --- | --- | --- | --- | --- | --- | --- | --- | --- | --- | --- | --- |
| All | Lean | HFD | All | Lean | HFD | All | Lean | HFD | All | Lean | HFD | All | HFD |
| $\dot{V}_E$ | EIP + Exp. T | BPM | $\dot{V}_E$ | $\dot{V}_E$ | BPM | Vol/EE | EIP + Exp. T | Vol/EE | $\dot{V}_E$ | $\dot{V}_E$ | $\dot{V}_E$ | $\dot{V}_E$ | BPM |
| -0.59*** | 0.67*** | -0.60** | -0.54** | -0.42* | -0.60** | -0.57*** | 0.64** | -0.57** | -0.48** | -0.48* | -0.54** | -0.40** | -0.61** |
| Vol/EE | EIP | $\dot{V}_E$ | BPM | EIP + Exp. T | $\dot{V}_E$ | $\dot{V}_E$ | EIP | $\dot{V}_E$ | Vol/EE | Vol/EE | Vol/EE | BPM | $\dot{V}_E$ |
| -0.59*** | 0.64** | -0.60** | -0.42** | 0.40* | -0.54** | -0.53*** | 0.59** | -0.54** | -0.46** | -0.45* | -0.47* | -0.39** | -0.57** |
| BPM | Vol/EE | Vol/EE | Vol/EE | BPM | Insp. T | EIP + Exp. T | Vol/EE | BPM | BPM | BPM | Insp. Time | EIP + Exp. T | EIP + Exp. T |
| -0.53*** | -0.62** | -0.56** | -0.41** | -0.40* | 0.53** | 0.51*** | -0.56** | -0.50* | -0.41** | -0.44* | 0.46* | 0.38** | 0.56** |
| EIP | $\dot{V}_E$ | Insp. Time | EIP + Exp. T | Insp. Time | EIP + Exp. T | BPM | BPM | Insp. T | EIP + Exp. T | EIP + Exp. T | EIP + Exp. T | Insp. T | Insp. T |
| 0.51*** | -0.62** | 0.55** | 0.39** | 0.39* | 0.53** | -0.49*** | -0.56** | 0.47* | 0.36* | 0.42* | 0.41* | 0.34* | 0.54** |
| EIP + Exp. T | BPM | EIP + Exp. T | EIP | Vol/EE | Vol/EE | EIP | $\dot{V}_E$ | Volume | Insp. T | Insp. T | - | EIP | Vol/EE |
| 0.51*** | -0.60** | 0.44* | 0.36* | -0.38* | -0.47* | 0.49*** | -0.54** | -0.45* | 0.35* | 0.42* | - | 0.31* | -0.51** |
| Volume | Insp. T | EF50 | Insp. T | - | EIP | EF50 | Insp. Time | EF50 | Volume | EIP | - | EF50 | EIP |
| -0.44** | 0.54** | -0.41* | 0.35* | - | 0.44* | -0.42** | 0.48* | -0.44* | -0.36* | 0.40* | - | -0.27* | 0.43* |
| EF50 | Volume | EIP | Volume | - | - | Insp. T | EF50 | EIP + Exp. T | EIP | EF50 | - | - | - |
| -0.41** | -0.49* | 0.37* | -0.29* | - | - | 0.42** | -0.45* | 0.36* | 0.34* | -0.38* | - | - | - |
| Insp. T | EF50 | Volume | EF50 | - | - | Volume | - | - | EF50 | - | - | - | - |
| 0.45** | -0.49* | -0.36* | -0.28* | - | - | -0.40** | - | - | -0.30* | - | - | - | - |

\*P<0.05; \*\*P<0.005; \*\*\*P<0.0001 listed next to r value; Breaths per Minute: BPM; End-Inspiratory Pause: EIP; End-Inspiratory Pause +
Expiratory Time: EIP + Exp. T; Inspiratory Time: Insp. T; Mid-Expiratory Flow: EF50; Minute Ventilation: $\dot{V}_E$ ; Volume Per Energy Expenditure:
Vol/EE

**Table 2.** Linear Regression Equations Comparing Maximal Methacholine Response Measured via FlexiVent and Head-Out Plethysmography.

|  |  |  |  |  |
| --- | --- | --- | --- | --- |
| FlexiVent Variable | Diet | Equation | P Value | R<sup>2</sup> |
| Max(Rrs) | All | $= -(9.85 \times \text{Vol}) - (0.01 \times \text{BPM}) + (3.73 \times \text{EIP}) - 0.19$ | <0.0001 | 0.416 |
| | Chow | $= -(0.33 \times \text{EF50}) + (7.02 \times \text{EIP}) - 0.01$ | <0.0001 | 0.531 |
| | HFD | $= -(0.01 \times \text{BPM}) + 0.19$ | 0.0004 | 0.361 |
| Max(Ers) | All | $= -(0.19 \times \text{BPM}) + 4.18$ | 0.001 | 0.174 |
| | Chow | $= -(0.46 \times \dot{V}_E) - 0.05$ | 0.027 | 0.178 |
| | HFD | $= -(4.16 \times \dot{V}_E) + (0.0 \times \text{Vol/EE}) - 11.11$ | 0.001 | 0.388 |
| Max(Rn) | All | $= (2.39 \times \text{EIP}) - (0.0001 \times \text{Vol/EE}) - 0.06$ | < 0.0001 | 0.357 |
| | Chow | $= -(0.23 \times \text{EF50}) + (4.90 \times \text{EIP}) - 0.09$ | 0.0005 | 0.460 |
| | HFD | $= -(6.92 \times \text{Vol}) + (10.44 \times \text{IT}) - 0.12$ | 0.001 | 0.396 |
| Max(G) | All | $= -(51.77 \times \text{Vol}) - (0.03 \times \text{BPM}) - 0.84$ | 0.0005 | 0.242 |
| | Chow | $= -(0.12 \times \dot{V}_E) + 0.11$ | 0.010 | 0.232 |
| | HFD | $= -(0.08 \times \text{BPM}) + 0.57$ | 0.003 | 0.272 |
| Max(H) | All | $= -(0.10 \times \text{BPM}) + 1.83$ | 0.003 | 0.150 |
|  | Chow |  |  |  |
| | HFD | $= -(0.22 \times \text{BPM}) - 2.10$ | 0.0003 | 0.377 |
| Breaths Per Minute: BPM; End-Inspiratory Pause: EIP; Volume: Vol; Inspiratory Time: IT; Mid-Expiratory Flow: EF50; Inspiratory Minute Ventilation: $\dot{V}_E$; Volume Per Energy Expenditure: Vol/EE | | | | |

**Figure 4.**
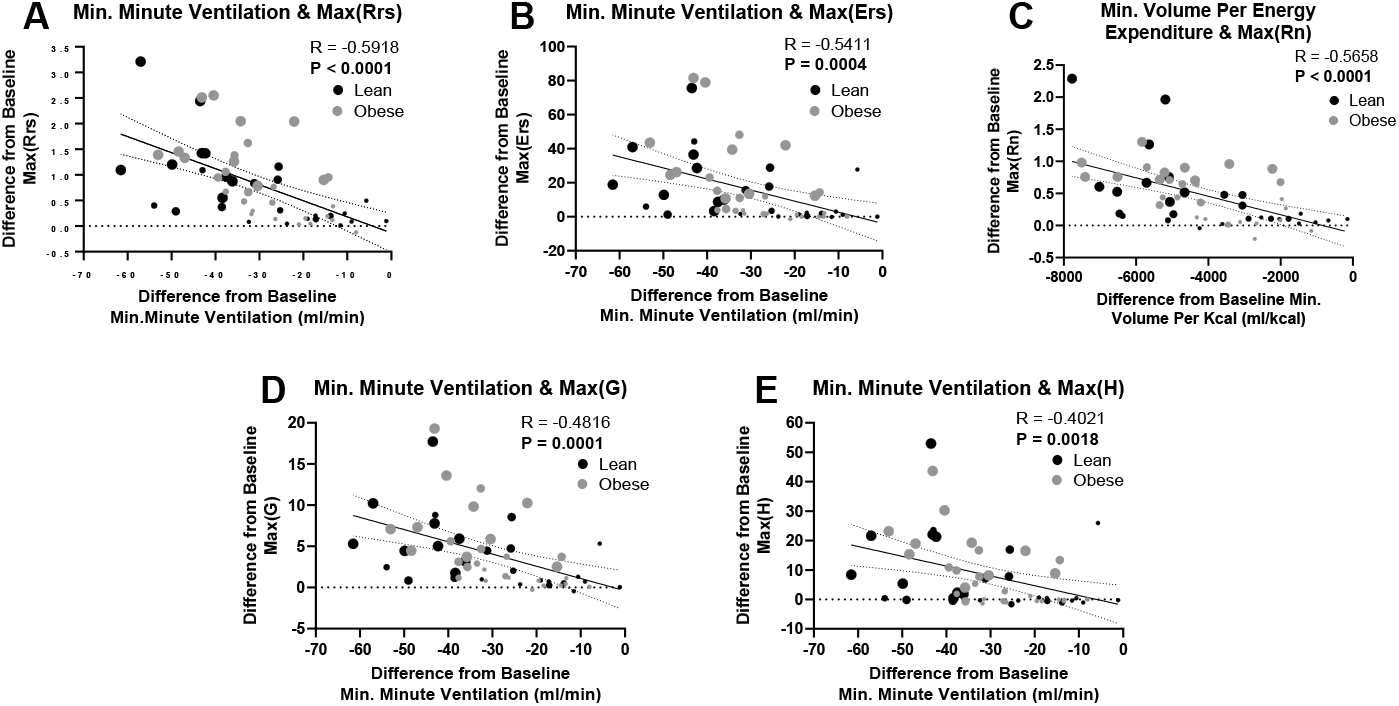
Most significant correlations between variables collected via head-out plethysmography to variables collected with the flexiVent system in response to methacholine. Results presented as maximal change from baseline for each variable in response to each methacholine exposure (0, 25, 50, 100 mg/ml at flow rate of 0.2 ml/30sec head-out plethysmography or flow rate 0.07ml/10sec forced oscillation technique) A) Maximal airway resistance, Max(Rrs), B) Maximal airway elastance, Max(Ers), C) Maximal resistance of conducting airways, Max(Rn), D) Maximal tissue damping, Max(G), and E) Maximal tissue elastance, Max(H). Solid line=linear regression best fit; dotted line=95% confidence interval. Each male mouse is represented by 4 data points representing the 4 doses of methacholine and the size of each dot represents methacholine dose (smallest dot = 0 mg/ml, largest dot = 100 mg/ml). Doses 0 & 100 mg/ml lean n=9 obese n=10; doses 25 & 50 mg/ml obese n=5 lean n=6. Analyzed with Pearson’s correlation coefficient.

To investigate how obesity may affect the correlations between variables measured by head-out plethysmography and forced oscillation technique, we repeated the correlation analysis separately for chow fed and diet-induced obese mice. In lean mice end-inspiratory pause + expiratory time (time of pause occurring at end of inspiration and the time of expiration) was most significantly associated with Max(Rrs) and Max(Rn) (r=0.67 P<0.0001, and r=0.64 P=0.0002 respectively) and 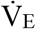 was most significantly associated with Max(Ers) and Max(G) (r=-0.42 P=0.027, and r=-0.48 P=0.0095, respectively; Online Data Supplement Figure E1). In obese mice, breaths per minute was most significantly associated with Max(Rrs), Max(Ers), and Max(H) (r=-0.60 P=0.0004, r=-0.60 P=0.0004, and r=-0.61 P=0.003 respectively; Online Data Supplement Figure E2). 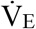 was most significantly associated with Max(G) (r=-0.54 P=0.003) and volume accounting for energy expenditure was most significantly associated with Max(Rn) (r=-0.57 P=0.001; Online Data Supplement Figure E2). The variables that remained in the regression equations after stepwise removal of variables with P > 0.10 differed by diet (Table 2).

### Role of Hepatic GABA Production & the Hepatic Vagal Afferent Nerve

To assess if obesity-associated alterations in breathing are driven by hepatic GABA production, we treated diet-induced obese male mice for four weeks with an ASO targeted to GABA-Transaminase (ABAT ASO) or a scrambled control (Control) and assessed tidal breathing after treatment using the head-out plethysmography system. This treatment paradigm reduces GABA-Transaminase mRNA expression by >98% and eliminates the obesity-induced rise in ex vivo liver slice GABA release(19, 22). Knocking down GABA-Transaminase did not alter tidal breathing (Figure 5) for any variable assessed.

**Figure 5.**
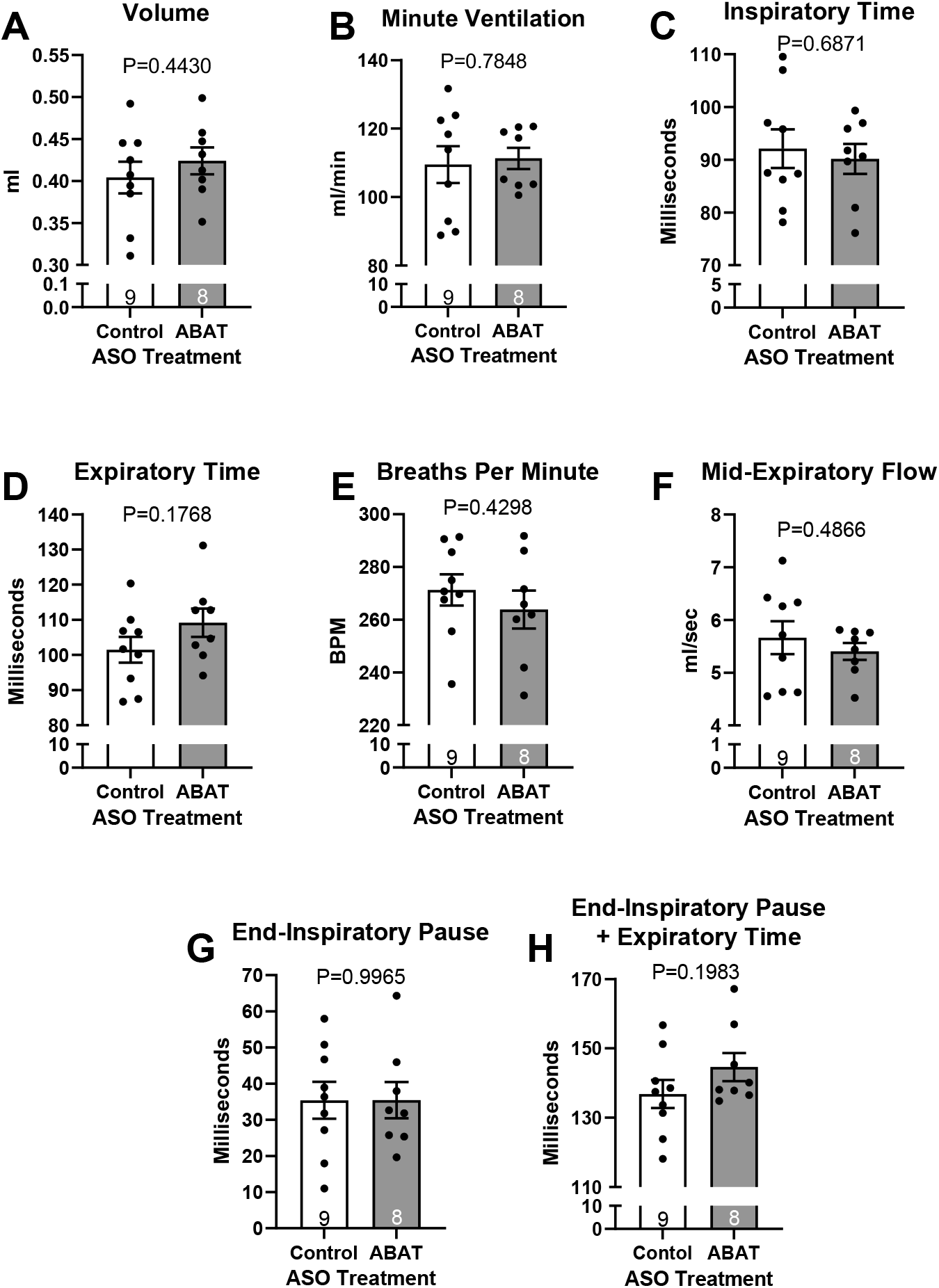
Tidal breathing assessed with head-out plethysmography in diet-induced obese male mice treated for 4 weeks with an antisense oligonucleotide targeted to GABA-T (ABAT ASO) or scrambled control (Control ASO). ABAT ASO decreases hepatic GABA production. Results presented as mean + SEM; analyzed with Students T-Test.

To understand if the hepatic vagal nerve was involved in the obesity-associated declines in lung function, we severed the hepatic vagal nerve near the esophagus. Hepatic vagotomy did not affect breathing in lean, chow-fed mice or in obese mice after 3, 6, and 9 weeks on a high fat diet (Figure 6).

**Figure 6.**
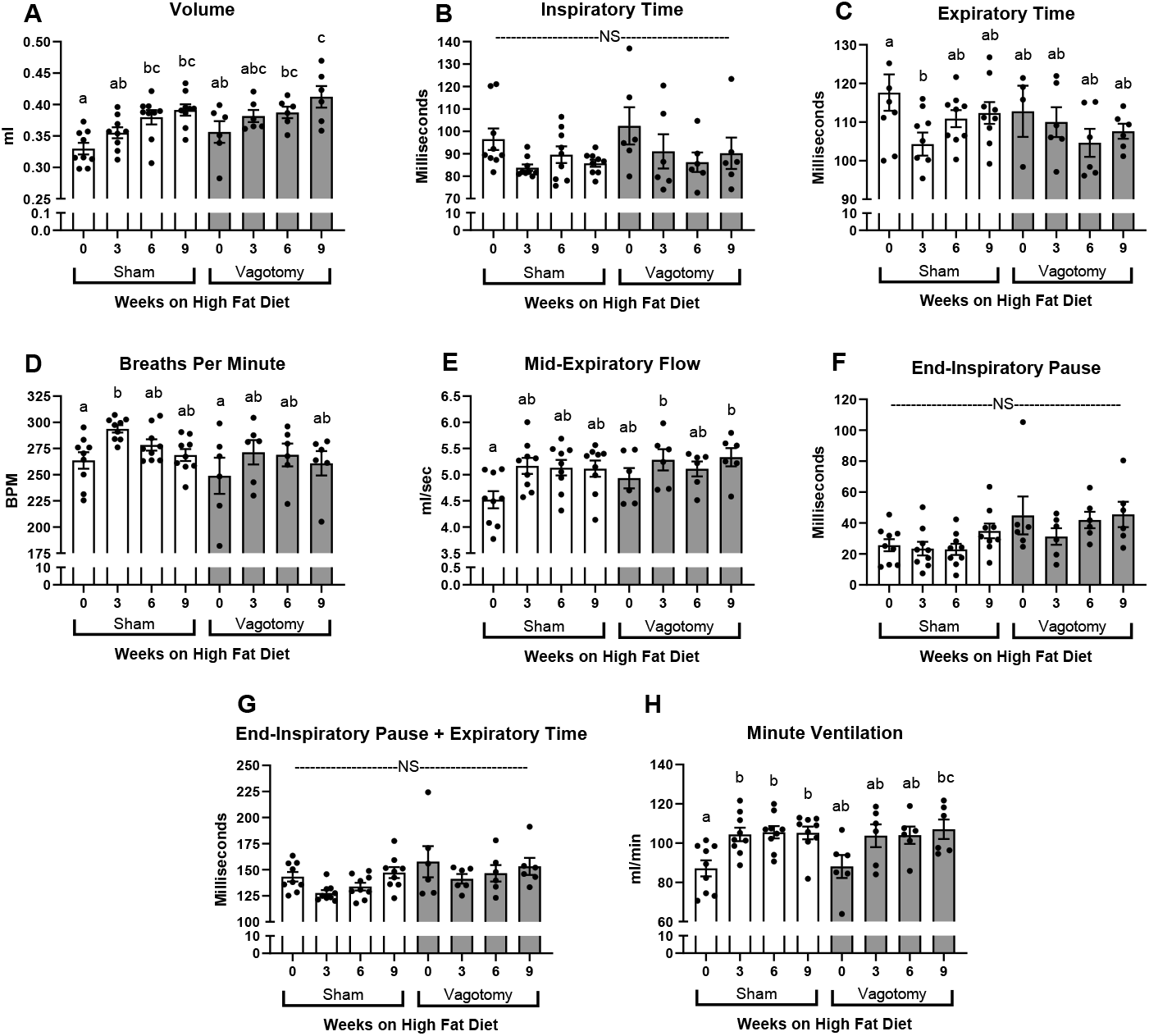
Tidal breathing after 0, 3, 6 and 9 weeks on a high fat diet in male mice after undergoing either a hepatic vagotomy (severing the hepatic afferent vagal nerve) or sham surgery. NS=no significance different letters signify statistically significant difference analyzed with mixed model ANOVA; results presented as mean + SEM sham surgery n=9, vagotomy surgery n=6.

## Discussion

Head-out plethysmography may provide insight into regulation of pulmonary function that is not available using forced oscillation technique. We measured obesity-associated alterations in alters tidal breathing after 9 weeks on a high fat diet in mice, yet, observed no obesity-induced differences in lung mechanics (Figures 1 & 3). Similar to observations in humans, we showed obesity increased minute ventilation (Figure 1C)(23). Our results recapitulate the findings from the leptin deficient (ob/ob) mouse in which minute ventilation is also increased further validating our results(24).

In humans, only morbid obesity is associated with a decrease in tidal volume as compared to healthy weight controls; overweight and obesity are not associated with decreased tidal volume(25, 26). Individuals with obesity also have decreased total lung capacity, functional residual capacity, and expiratory reserve volume(23, 25). In contrast, we showed that obesity increased tidal volume in diet-induced obese mice (Figure 1A). Tidal volume measured in genetically obese, ob/ob or Cpe^fat^, does not differ from lean wild-type controls(27, 28).

Unlike volume and minute ventilation, mid-expiratory flow is not commonly reported in human studies. Instead, the flow of a forceful exhalation (forced expiratory volume, forced expiratory volume over 1 second, ratio of forced expiratory volume over one second divided by total forced expiratory volume, or maximal expiratory flow at 50% volume) is assessed, thus, direct comparison with tidal breathing studies in rodents is difficult. Forceful expiratory flow is decreased in humans with obesity(26). Diet-induced obesity increased mid-expiratory flow consistent with our reported increase in volume (Figure 1G).

Obesity increases breaths per minute and decreases expiratory time in humans(23, 25). Although breaths per minute was not affected by diet-induced obesity in mice, we did observe a decrease in expiratory time (P < 0.0001; Figure 1E). This decrease in expiratory time was offset by an increase in end-inspiratory pause (P = 0.0091; Figure 1H). Also referred to as braking, this measure acutely increases in response to stimulation of the vagal nerve ending or a pulmonary irritant at the alveolar level(29). The increase in end-inspiratory pause is a measure specific to small animals because their chest wall is more flexible than humans(30). Extending the end-inspiratory pause helps to get air further into the lung(31). Given that end-inspiratory pause is specific to small animals, our measured decrease in expiratory time would be measured as an increase in breaths per minute in humans.

Though we showed that obesity altered tidal breathing, obesity did not induce changes in the response to methacholine as assessed by either head-out plethysmography or forced oscillation technique (Figures 2 and 3). Conversely, obesity has been reported to increase the response to methacholine in humans(11, 32). However, obesity-associated increases in methacholine response may be a result of decreases in lung volume(11). Persistent obesity to induce structural changes to the airway may instead be necessary for the measured increase in hyperresponsiveness. In fact, methacholine response was not affected after 9-12 weeks of high fat diet feeding in mice, but high fat diet feeding for >15 weeks results in changes to airway mechanics at baseline and in response to methacholine(33, 34).

While other studies have compared the response to methacholine using a non-invasive technique with the response to methacholine measured by forced oscillation technique; we are providing the most comprehensive comparison of variables measured using two systems in the same mice(35–38). We are also the first to show that these associations are affected by diet (Tables 1 & 2, Online Data Supplement Figures E1 & E2).

Obesity is closely associated with hepatic lipid accumulation, in fact, pulmonary function is inversely associated with the degree of hepatic steatosis(15, 39). We hypothesized that hepatic GABA production, associated with increased hepatic lipids, and subsequent altered firing of the hepatic vagal afferent nerve may mediate obesity-induced changes in lung function. Neither eliminating hepatic GABA production nor severing the hepatic vagal afferent nerve affected tidal breathing in diet induced obese mice (Figures 5 & 6). Still, with the recognition that the liver is an endocrine organ, there remain plenty of potential signaling molecules by which hepatic lipid accumulation alters lung function(40–42). Additionally, several non-hepatocentric mechanisms including increased circulating leptin, inflammation, and/or mechanical factors may mediate the obesity-induced changes in pulmonary function(43–45).

Herein we have shown that a leak-free head-out plethysmography system can assess obesity-associated alterations in tidal breathing in mice and we have validated our system by comparing against the gold-standard, forced oscillation technique. Our variable-pressure head-out plethysmography system provides a useful method for future studies using a mouse model to investigate mechanisms inducing obesity-associated pulmonary dysfunction.

## Acknowledgements

The authors wish to thank Dr. Frank Duca for allowing us to use their indirect calorimetry machine and ECHO MRI.

## Author Contributions^1^

Stephanie Bruggink: Completion of head-out plethysmography measurements, development of head-out plethysmography system, data interpretation, and manuscript preparation

William Pederson: Completion of forced oscillation technique measurements, & analysis, manuscript preparation

Kyle Kentch: Development of head-out plethysmography system and software

Jason Kronenfeld: Analysis of data collected with head-out plethysmography system

Julie Ledford: Guidance on forced oscillation technique measures and manuscript preparation

Benjamin Renquist: Development of head-out plethysmography system, statistical analysis, data interpretation, and manuscript preparation

## Abbreviations

ASO: Antisense Oligonucleotide
BPM: Breaths per Minute
EIP: End-Inspiratory Pause
GABA: Gamma Aminobutyric Acid
GABA-T: GABA-Transaminase
HVAN Hepatic: Vagal Afferent Nerve
Max(Ers): Maximal Airway Elastance
Max(Rrs): Maximal Airway Resistance
Max(Rn): Maximal Resistance of Conducting Airways
Max(G): Maximal Tissue Damping
Max(H): Maximal Tissue Elastance
EF50: Mid-Expiratory Flow
Minute Ventilation: 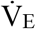

## Footnotes

1 Funding Sources: ABRC ADHS14-082986 (BJR), ABRC ADHS17-00002043 (BJR), T32 HL 007249 (SB), ARCS-Phoenix Chapter (SB)

