## Supplemental Material for "Obesity-Associated Alterations in Lung Function in Mice Measured with Head-Out Plethysmography"

**Online Data Supplement**

**Materials & Methods**

*Mice*

C57BL/6J male mice were bred in-house with lines originating from Jackson Laboratories (Bar harbor, ME). Lean mice were fed chow diet (Tekland WI 7013 NIH-31, 3.1 kcal/g, kcal from fat 18%, carbohydrate 59%, protein 23%). To induce obesity, at 12 weeks of age mice were provided a high-fat diet for 9 weeks (Tekland WI TD 06414, 5.1 kcal/g, kcal from fat 60.3%, carbohydrate 21.4%, protein 18.3%). We used male mice for all studies because of their propensity to develop obesity. Mice were housed 3-5 per cage and housed in a 14-hour light/10-hour dark schedule. Weights for all studies are reported in Online Data Supplement Table E1. All data graphed in GraphPad Prism software (GraphPad Software; San Diego, CA).

To compare lean and obese mice, we corrected volume for oxygen demand. Metabolic rate in lean and diet-induced obese mice was assessed using indirect calorimetry (Promethion Systems Sable Systems International, North Las Vegas, NV). Calorimetry occurred over 3 days with the first day serving as a day of acclimation and the final 2 days serving as the days of measurement.

*Hepatic Vagotomy*

12-week-old mice were randomized to sham or vagotomy surgery and surgery was completed as previously described (20). Briefly, mice were anesthetized under isoflurane, the abdomen was shaved and sterilized, and a midline incision was made. The vagus was identified as it branched from the esophagus. In vagotomized mice the hepatic vagal nerve was severed while it was left intact in sham operated mice. The incision was closed, and mice given a slow-release buprenorphine analgesic (1.2 mg/kg slow release, subcutaneous). Food intake and body weight was measured for 7 days post-surgery and lung function was first assessed after mice had recovered to pre-surgical weight (week 0 on HFD) and subsequently 3, 6, and 9 weeks after being place on a HFD.

*Antisense Oligonucleotide Treatment*

Obese mice were treated for four weeks with an antisense oligonucleotide (ASO; IP 12.5 mg/kg biweekly injection) as previously described(19). Mice were randomized to either a scrambled control ASO (IONIS 549144; 5′- GGCCAATACGCCGTCA-3′; Ionis Pharmaceuticals Carlsbad, CA) or an ASO targeting the enzyme GABA-Transaminase (ABAT ASO; IONIS 1160575; 5′- AAGCTATGGACTCGGT-3′; Ionis Pharmaceuticals Carlsbad, CA). ABAT ASO treatment is effective in knocking-down hepatic GABA-transaminase mRNA expression by >98% and reduce hepatic GABA production(19). After four weeks, tidal breathing was assessed using our head-out plethysmography system.

*Comparison of Breathing Measures*

For all studies mice were exposed to increasing concentrations of methacholine (MP Biomedicals LLC; MP 220). For head-out plethysmography measures, we delivered a total of 0, 5, 10, and 20 mg aerosolized onto the face of the mouse by exposing mice to 0, 25, 50, 100 mg/ml in sterile PBS at a flow rate of 0.2ml/30sec. We compared this measure to the response to methacholine collected using forced oscillation technique. We delivered a total of 0.21, 1.75, 3.5, and 7 mg by administering methacholine intratracheally at concentrations of 3, 25, 50, 100 mg/ml in sterile PBS at a flow 0.07ml/10sec. We used the 3 mg/ml dose as a 0 dose in comparison studies. For comparison studies each mouse is represented by 4 data points representing the 4 doses of methacholine.

| **Online Data Supplement Table E1. Average Mouse Weight for Each Study** | | |
| --- | --- | --- |
| **Study** | **Average Mouse Weight ± SEM (grams)** | |
| Lean Dose Response | 24.4 ± 0.7 | |
| Obese Dose Response | 40.8 ± 1.2 | |
| Subset of Lean Dose Response used for Forced Oscillation Technique Measure | 26.9 ± 0.7 | |
| Subset of Obese Dose Response used for Forced Oscillation Technique Measure | 35.6 ± 0.8 | |
| Antisense Oligonucleotide Study | 45.8 ± 0.8 | |
| Sham & Vagotomy Study Weeks on High-Fat Diet | Sham | Vagotomy |
| Week 0 | 27.1 ± 1.0 | 27.0 ± 0.6 |
| Week 3 | 33.2 ± 1.3 | 33.5 ± 1.0 |
| Week 6 | 36.9 ± 2.1 | 37.0 ± 1.5 |
| Week 9 | 40.0 ± 2.6 | 38.8 ± 2.1 |

**Supplementary Figure E1.** Most significant correlations in lean male mice between the variables collected via head-out plethysmography to variables collected using the flexiVent system in response to methacholine. Results presented maximal change from baseline for each variable in response to methacholine exposure (0, 25, 50, 100 mg/ml at flow rate of 0.2 ml exposed for 30 seconds for head-out plethysmography and 10 sec for forced oscillation technique). A) Maximal airway resistance, Max(Rrs), B) Maximal airway elastance, Max(Ers), C) Maximal resistance of conducting airways, Max(Rn), and D) Maximal tissue damping, Max(G). Solid line=linear regression best fit; dotted line=95% confidence interval. Each mouse is represented by 4 data points representing the 4 doses of methacholine and the size of each dot represents methacholine dose (smallest dot = 0 mg/ml, largest dot = 100 mg/ml). Doses 0 & 100 mg/ml n=9; doses 25 & 50 n=5. Analyzed with Pearson’s correlation coefficient.

**Supplementary Figure E2.** Most significant correlations in obese male mice between the variables collected via head-out plethysmography to variables collected using the flexiVent system in response to methacholine. Results presented as maximal change from baseline for each variable in response to methacholine exposure (0, 25, 50, 100 mg/ml at flow rate of 0.2 ml exposure 30 sec for head-out plethysmography and 10 sec for forced oscillation technique) A) Maximal airway resistance, Max(Rrs), B) Maximal airway elastance, Max(Ers), C) Maximal resistance of conducting airways, Max(Rn), D) Maximal tissue damping, Max(G), and E) Maximal tissue elastance, Max(H). Solid line=linear regression best fit; dotted line=95% confidence interval. Each mouse is represented by 4 data points representing the 4 doses of methacholine and the size of each dot represents methacholine dose (smallest dot = 0 mg/ml, largest dot = 100 mg/ml). Doses 0 & 100 mg/ml n=10; dose 25 n=4; dose 50 n=6. Analyzed with Pearson’s correlation coefficient.

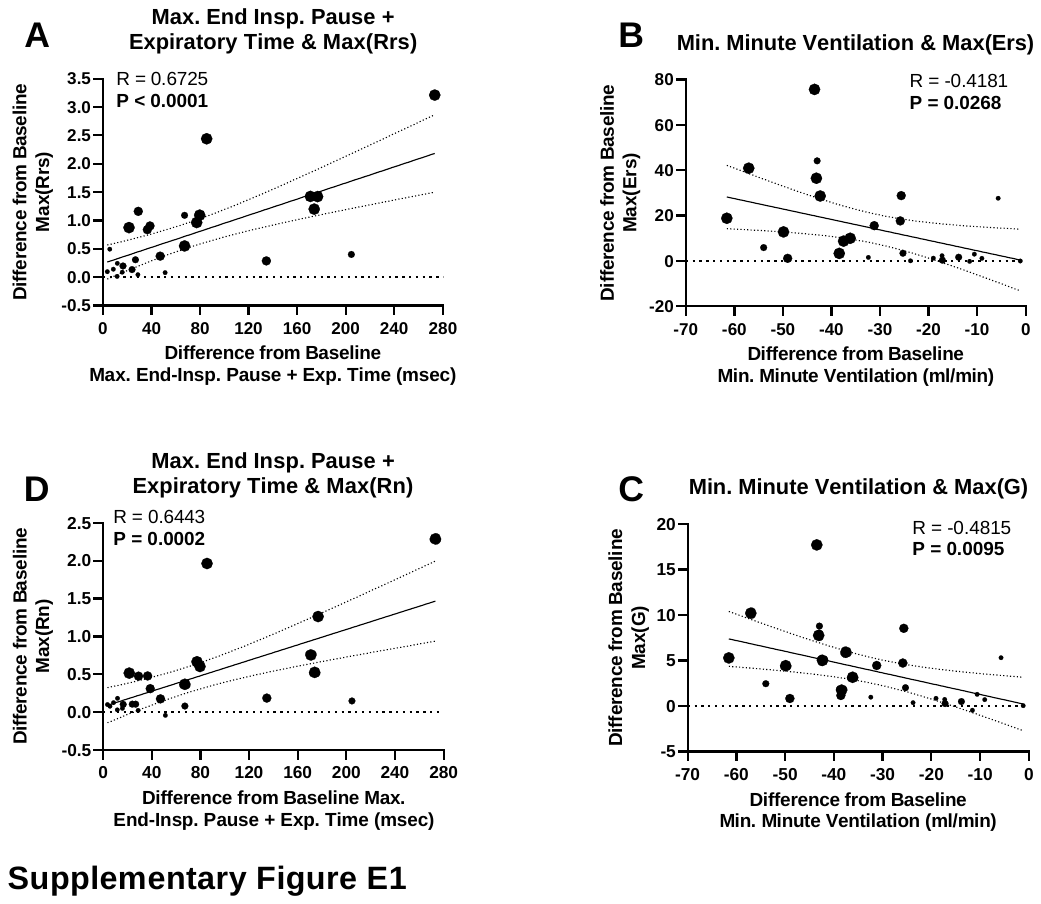

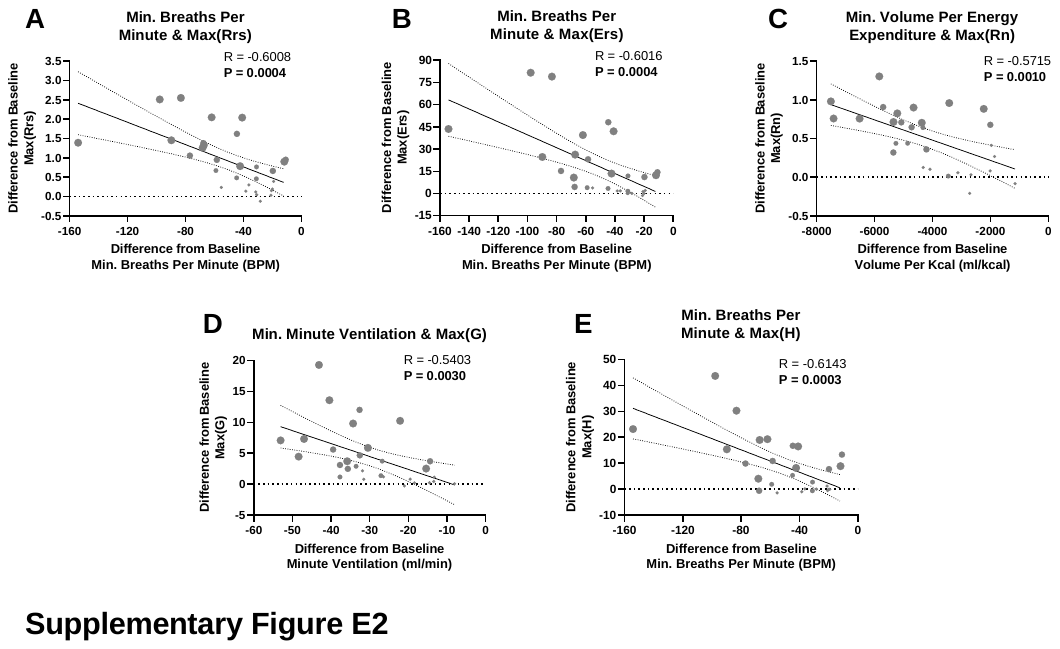
